# Fine-mapping cellular QTLs with RASQUAL and ATAC-seq

**DOI:** 10.1101/018788

**Authors:** Natsuhiko Kumasaka, Andrew Knights, Daniel Gaffney

## Abstract

When cellular traits are measured using high-throughput DNA sequencing quantitative trait loci (QTLs) manifest at two levels: population level differences between individuals and allelic differences between *cis*-haplotypes within individuals. We present RASQUAL (Robust Allele Specific QUAntitation and quality controL), a novel statistical approach for association mapping that integrates genetic effects and robust modelling of biases in next generation sequencing (NGS) data within a single, probabilistic framework. RASQUAL substantially improves causal variant localisation and sensitivity of association detection over existing methods in RNA-seq, DNaseI-seq and ChIP-seq data. We illustrate how RASQUAL can be used to maximise association detection by generating the first map of chromatin accessibility QTLs (caQTLs) in a European population using ATAC-seq. Despite a modest sample size, we identified 2,706 independent caQTLs (FDR 10%) and illustrate how RASQUAL’s improved causal variant localisation provides powerful information for fine-mapping disease-associated variants. We also map “multipeak” caQTLs, identical genetic associations found across multiple, independent open chromatin regions and illustrate how genetic signals in ATAC-seq data can be used to link distal regulatory elements with gene promoters. Our results highlight how joint modelling of population and allele-specific genetic signals can improve functional interpretation of noncoding variation.

## Introduction

Association mapping of cellular traits is a powerful approach for understanding the function of genetic variation. Cellular traits that can be quantified by next generation sequencing (NGS) are particularly useful for association analysis because they provide highly quantitative information about the phenotype of interest and can easily be scaled genome-wide. Population scale studies using NGS-based cell phenotypes such as RNA-seq, ChIP-seq and DNaseI-seq have revealed an abundance QTLs for gene expression and isoform abundance [1–4], chromatin accessibility [5], histone modification, transcription factor binding (TF) [6–9] and DNA methylation [10], providing precise molecular information on the functions of human genetic variation at high resolution. However the effect sizes of many common variants are modest meaning that association analysis typically requires large sample sizes, which can be problematic when assays are labour intensive or cellular material is difficult to obtain. Furthermore, even well-powered studies can struggle to accurately fine-map causal variants.

A significant advantage of NGS-based cell phenotyping is the ability to identify allelespecific (AS) differences in traits between maternal and paternal chromosomes [11]. AS differences can arise when a sequenced individual is heterozygous for a *cis*-acting causal variant and several studies have highlighted abundant AS changes in a variety of cell traits [1, 2, 5, 7]. Uniquely, AS signals provide information both about the existence of a QTL and about the identity of the causal variant [12] in a single sample. However, although both population and AS signals provide complementary information about genetic associations, principled approaches for combining them are lacking. In part this is because AS signals are challenging to analyse: allelic imbalance (AI) can also be produced by a wide variety of technical factors including reference mapping bias [13], the presence of collapsed repeats [14], PCR amplification bias [15, 16] and sequencing error [17]. Biological phenomena such as imprinting or random allelic inactivation (RAI) [6, 15] can also produce AI when no *cis*-QTL exists. Genotyping errors can also be a serious problem for AS analysis, particularly in cases where homozygous SNPs located within a sequenced feature (feature SNPs, fSNPs) are miscalled as heterozygous [6]. Effective use of AS information must take account of these biases to avoid high false positive rates [15]. Previous strategies to address these problems have included the creation of personal reference genomes for read mapping, read masking, genomic blacklists or simulation strategies to compute genome-wide mapping probabilities that account for reference bias effects. However, it is challenging to set sensible values for the thresholds that these strategies rely on: overly conservative settings can lead to a loss of power while overly liberal settings may inflate the false positive rate. Additionally, genome wide simulations, custom read filtering and alignment steps significantly increase the time and computational burden required for analysis.

Here we describe a novel statistical method, RASQUAL (Robust Allele Specific Quantitation and quality controL), that integrates population level changes, AS signals and technical biases on NGS-based cell phenotypes into a single, probabilistic framework for association mapping. RASQUAL can be applied to existing NGS data sets without requiring data filtering, masking or the creation of personalised reference genomes. When applied to RNA-seq, ChIP-seq and DNaseI-seq data sets, RASQUAL significantly outperformed existing methods, both in its ability to detect QTLs and to fine-map putatively causal variants. We used RASQUAL to generate the first map of chromatin accessibility QTLs in a European population using ATAC-seq [18]). Despite a modest sample size of 24 individuals, RASQUAL detected over 2700 independent chromatin accessibility QTLs (FDR 10%) providing a rich resource for the functional interpretation of human noncoding variation.

## Results

### Rationale

To illustrate the strategy employed by RASQUAL, consider a sequenced feature such as a ChIP-seq peak or the union of exons across a transcript, that is affected by a single *cis*-regulatory SNP (rSNP) such that the total number of reads mapped onto the feature correlates with rSNP genotype (the population signal; Fig. 1a). When sequenced reads overlap SNPs located inside a sequenced feature (fSNPs), AS differences can be detected by comparing the numbers of reads that map to one or other allele of the fSNP (the AS signal; Fig. 1a). Combining these signals achieves two objectives in QTL mapping (i) identification of the rSNP whose genotypes correlates with the total fragment count and (ii) confirmation of AI at one or more fSNPs in individuals that are heterozygous for the rSNP (Fig. 1b).

**Figure 1:**
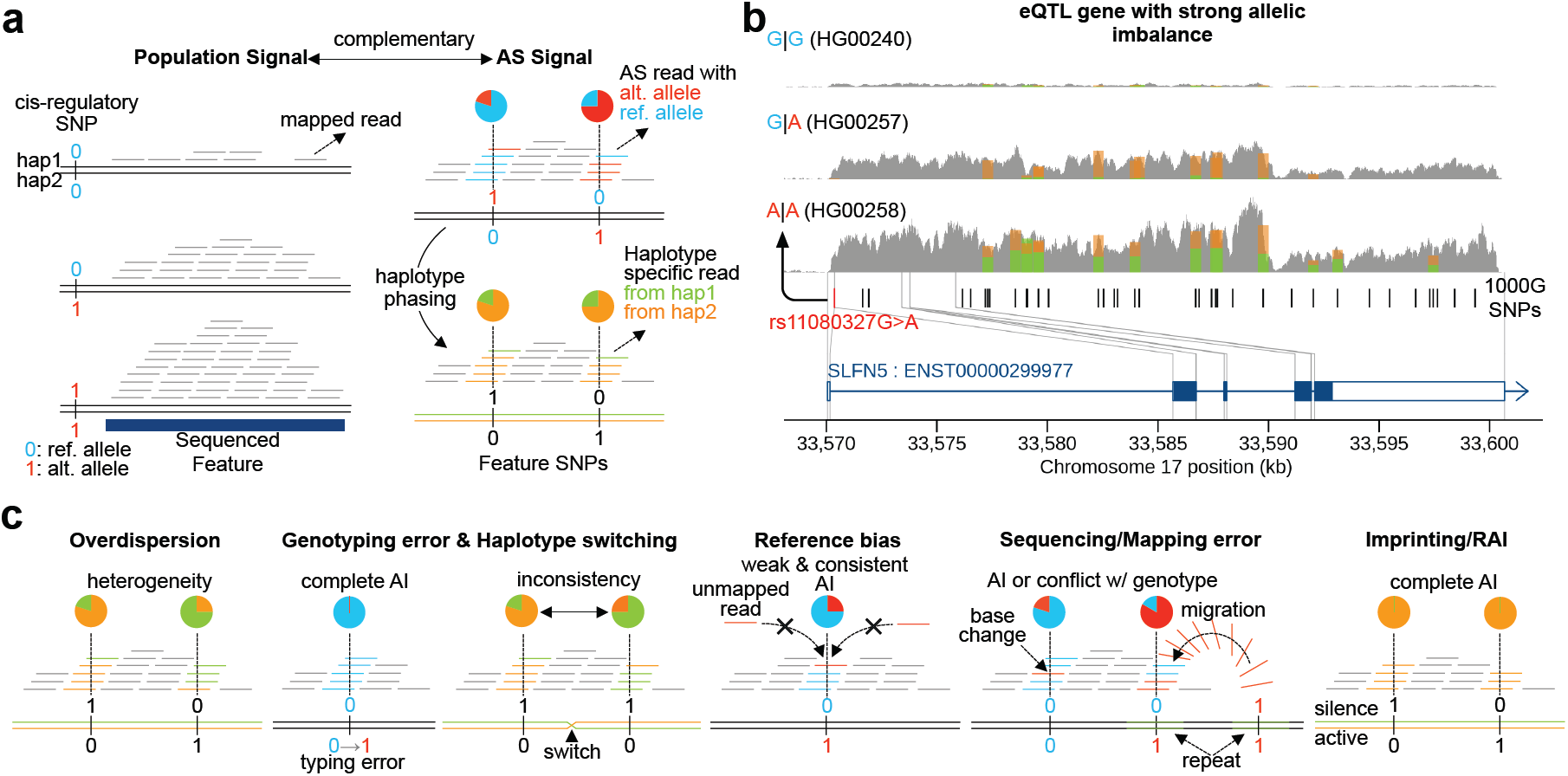
Schematic of RASQUAL approach. Throughout, reference and alternate alleles are coloured blue and red and coded 0 or 1, respectively, while alternative haplotype are coloured orange and green, respectively. (a) Plot illustrates the two sources of input data to RASQUAL: population and AS signals, as observed from NGS data. Right hand panel illustrates conversion between allele and haplotype specific expression. (b) Spliced coverage plot of a gene (*SLFN*5) with a strong eQTL in three individuals with three different genotypes at the predicted rSNP (rs11080327G*>*A). The stacked bars indicate haplotype-specific fragment counts at heterozygous fSNPs. Here, the alternative allele “A” up-regulates the expression level (the population signal) and strong AI is observed in the heterozygous individual (HG00257) (AS signal) in which the haplotype 2 (orange) is linked to the alternative allele. (c) Visual representation of the key RASQUAL features and parameters.

Two statistical approaches have previously been proposed to combine population and AS signals for QTL mapping: TReCASE [19] and the combined haplotype test (CHT) [6,19]. RASQUAL improves on these methods via (i) robust estimation of read count overdispersion for both between-individual feature counts and within-individual AS counts, (ii) iterative correction of genotyping errors and haplotype switching between rSNP and each fSNP and (iii) explicit modelling of a broad range of technical biases in AS data by using information from all individuals (Fig. 1c; Supplementary Table 1). A key novelty of our approach is the use of read counts at both heterozygous and homozygous fSNPs to significantly improve genotype error correction and the estimation of bias parameters (see Supplementary Methods for details). Parameter estimation and genotypes are iteratively updated during model fitting by an expectation-maximisation (EM) algorithm [20] to arrive at the final QTL call for each sequenced feature (Supplementary Fig. 1). This iterative approach also enables true QTLs to be distinguished from imprinting or RAI. For each feature, RASQUAL outputs a likelihood ratio test statistic for the hypothesis of a single QTL as well as estimated over-dispersion, reference allele mapping bias, sequencing/mapping error rate at each tested SNP and posterior probabilities for each genotype at the lead rSNP and fSNPs. RASQUAL also performs a separate likelihood ratio test for imprinting in the given feature. RASQUAL is entirely implemented in C with the software and accompanying documentation available from http://github.com/dg13/rasqual/.

### Combined AS and population signatures improves causal variant localisation

We first investigated the relative importance of the AS and population components of the RASQUAL model. We assessed power using an RNA-seq data set from 373 lymphoblastoid cell lines (LCLs) in European individuals generated by the gEUVADIS project [3]. Our analysis used a challenging test of model performance: how many QTLs mapped using the full data set could our model detect in a small subsample of the same data? We compared the numbers of eQTLs detected by RASQUAL in a subsample of 24 individuals of RNA-seq data with the set of “true positive” eQTLs provided by gEUVADIS project (see Online Methods). Our results clear show that RASQUAL’s combined allele-specific and population level information significantly outperformed either source alone with the joint model detecting, for example, 40% of eQTLs in the true positive set at false positive rate (FPR) at 10% compared with 32% and 29% for the population and allele-specific models (Fig. 2a, b).

**Figure 2:**
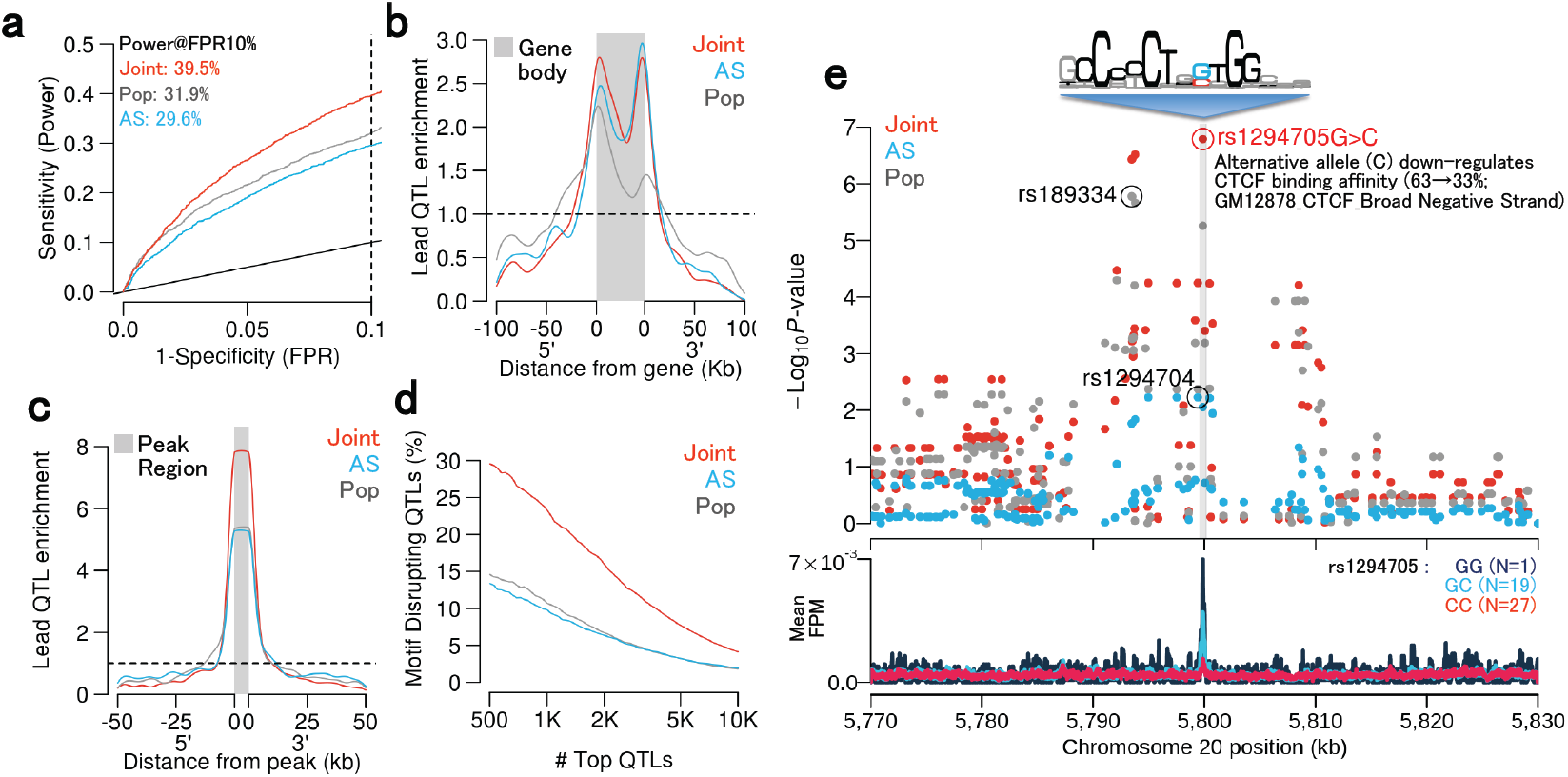
Comparing population-only, AS-only and combined models. In panels a-d, red curves indicate the joint RASQUAL model, blue indicates the AS-only signal and grey indicates the population-only signal. (a) ROC curves for detecting known eQTL genes (see Online Methods) for the three different models in a random subset of 24 individuals from gEUVADIS RNA-seq data [3]. Dotted line indicates FPR=10%. (b) Density plot shows the enrichment of top 1,000 lead eQTLs relative to the gene body and 5’/3’ flanking regions. (c) Density plot showing positional enrichment of the lead CTCF QTL SNPs near the CTCF peak, relative to all SNPs, aggregated over the top 1,000 detected CTCF QTLs. (d) The percentage of motif-disrupting lead SNPs in top *N* CTCF binding QTLs. Motif-disrupting SNPS were defined as SNPs located within a CTCF peak and putative CTCF motif, whose predicted allelic effect on binding, computed using CisBP position weight matrices [21], corresponded to an observed change in CTCF ChIP-seq peak height in the expected direction (see Online Methods). Ordering of the top QTLs was based on their statistical significance independently measured by the three models. (e) Regional plot of *P*-values around an example CTCF binding QTL (top panel) and CTCF ChIP-seq coverage plot stratified by the lead SNP detected by the joint model (rs1294705) (bottom panel). The sequencing logo (Accession M4325) was derived by the CisBP database analysis of ENCODE CTCF ChIP-seq for GM12878 conducted by Broad Institute.

Next we examined how RASQUAL’s combined model could improve the accuracy of fine-mapping. Here, we used a set of 47 ChiP-seq samples for CCCTC-binding factor (CTCF) in LCLs derived from European individuals [9]. The availability of population scale CTCF ChIP-seq data provided a unique opportunity to test fine mapping performance because causal CTCF QTLs are expected to frequently occur within a well-defined region: the relatively long and informative canonical CTCF binding motif. We defined a high confidence set of “motif-disrupting” putatively causal variants by identifying those SNPs that fulfilled three criteria: (i) they were located within CTCF peak regions (ii) they were located inside CTCF motif matches and (iii) there was concordance between the predicted and observed allelic effect on binding, where predicted allelic effects were computed using the CTCF position weight matrix from the CisBP database [21] (see Online Methods for details). RASQUAL’s combined model dramatically improved causal variant localisation. CTCF lead SNPs detected by a combined model were over twice as likely to be motif-disrupting: 29% of lead SNPs in our top 500 CTCF QTLs from the combined model occurred within the CTCF motif, compared with 14% and 13% of lead SNPs from the AS or population only models (Fig. 2c,d). An example of a putatively causal CTCF SNP that was successfully colocalised only by the combined model is shown in (Fig. 2e).

### RASQUAL outperforms existing methods for QTL detection and fine-mapping

We next compared RASQUAL with three other methods for QTL mapping in NGS-based traits: simple linear regression of log-transformed, principal component-corrected FPKM values, TRe-CASE, as implemented in the AS-seq package, and CHT as implemented in the WASP package [6]. A brief summary of the mathematical differences between TReCASE, CHT and RASQUAL is discussed in the Supplementary Methods and summarised in Supplementary Table 1. For this comparison, in addition to the RNA-seq and ChIP-seq data sets, we also analysed a data set of DNaseI-seq from 70 Yoruban individuals set [5] where we again compared QTLs detected in a subsample with a set of “true positive” DNase QTLs mapped using the full data (see Online Methods for details). Across all sample sizes in all data sets, RASQUAL significantly outperforms the other two methods (Fig. 3, Supplementary Fig. 3). At a false positive rate (FPR) of 10% RASQUAL detected between 50 and 130% more eQTLs and between 60 and 150% more DNase QTLs than simple linear regression and between 14 and 30% more eQTLs and between 9 and 24% more DNase QTLs than the next best performing method (Fig. 3a,b). Results for variant localisation were similarly improved with, for example, RASQUAL lead SNPs in the top 500 CTCF QTLs 2.5-fold more likely be “motif-disrupting” compared to simple linear regression and 50% more likely than next best performing method (Fig. 3c). In the majority of cases the next best performing method was CHT, although it performed significantly worse than both RASQUAL and TReCaSE for larger sample sizes. Across all data sets CHT also took significantly longer to run than RASQUAL, for example requiring 542 days of CPU time to analyse the CTCF ChIP-seq data set, compared with 36.2 CPU days for RASQUAL (Fig. 3d). Fixing the overdispersion parameter of CHT to the default value rather than estimating it from the data improved performance slightly for the eQTL data (Supplementary Figure 4), but hampered performance in the CTCF and DNase data, where very few QTLs were detected with the default overdispersion parameter. Analysis of permuted data showed that the RASQUAL model is well-calibrated under the null (Supplementary Figure 2).

**Figure 3:**
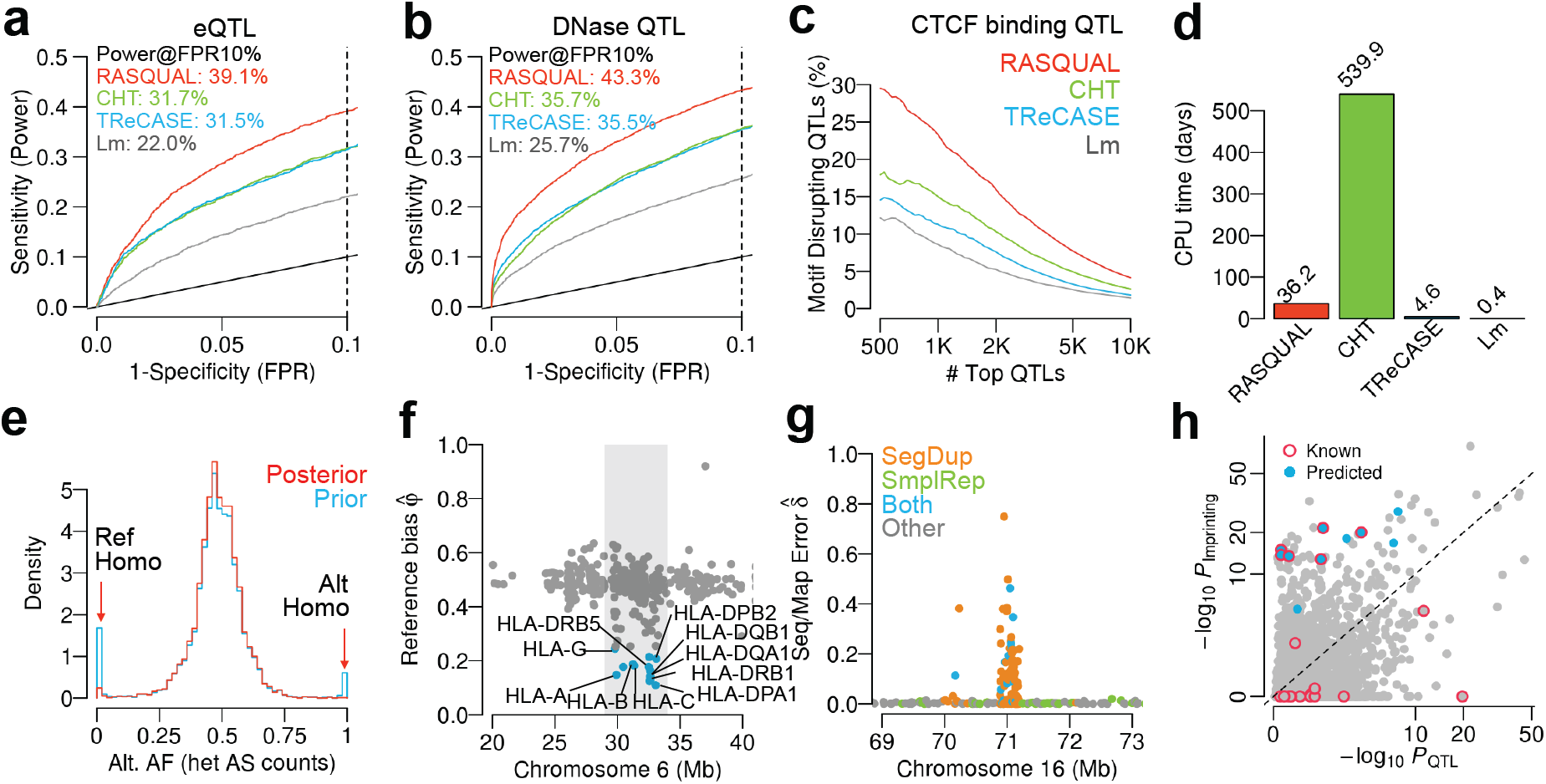
Comparison of RASQUAL with existing methods. RASQUAL was compared with the combined haplotype test (CHT), the TReCASE implemented in asSeq package (TReCASE) and with simple linear regression of log-transformed, principal component-corrected FPKM values (Lm). (a) ROC curves for detecting known eQTL genes (see Online Methods) in a random subset of 25 individuals from gEUVADIS RNA-seq data. Dotted line indicates FPR=10%. (b) ROC curves for detecting known DNaseI QTLs (see Online Methods) in a random subset of 25 individuals from DNaseI-seq data [5]. The dotted line indicates FPR=10%. (c) Percentage of motif-disrupting SNPs (see Online Methods) in top *N* lead CTCF-binding QTLs. Ordering of the top QTLs was based on their statistical significance independently measured by the four models. (d) CPU time in days required by each method to finish mapping CTCF QTLs genome-wide. (e) Spectrum of AI at putative heterozygous fSNPs (coverage depth *>* 20). Heterozygous fSNPs are called as maximum “a priori” genotype (blue) and maximum “a posteriori” genotype (red) (see Online Methods). (f) Genomic distribution of the reference bias parameter (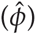) for the RNA-seq data estimated by RASQUAL around HLA region (chr6:28,477,797-33,448,354). The x-axis shows genomic position, the y-axis shows the reference bias parameter and each point corresponds to an individual gene. Genes with 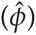 are coloured in blue and individual HLA Class I & II genes are labelled. (g) Example of a genomic distribution of the sequencing/mapping error 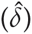 estimated by RASQUAL for the CTCF ChIP-seq data. Colours represent known segmental duplications (orange), simple repeats (green) or both (blue). (h) Comparison of *P*-values from the QTL model and the imprinting/RAI model of RASQUAL for the RNA-seq data. Blue points indicate predicted imprinted genes by RASQUAL and red circles indicate know imprinted genes from the Geneimprint website.

### Overdispersion and genotyping error

We next examined the ability of RASQUAL to handle two common features of NGS data that are problematic for AS analysis: read overdispersion and genotyping error. Although overdispersion of NGS data is well appreciated in the literature on differential expression (e.g. ref [22]), it is overlooked surprisingly often in AS analysis [23–29]. RASQUAL models overdispersion in total read counts and allele specific counts using a single parameter shared between the AS and population components of the model. Modelling overdispersion in this way provided a very substantial (2.6-fold) increase in power over a Poisson-binomial model (Supplementary Figure 5). One implication of this result is that using non-overdispersed distributions to model AS signals will significantly inflate the false positive rate, because random fluctuations in allelic ratios may not be properly accounted for (e.g. Supplementary Fig. 6a).

RASQUAL also employs a novel, iterative approach to genotyping error that refines imperfect genotype calls from genome imputation. Prior to model fitting, we observed an excess of heterozygous SNPs exhibiting complete monoallelic expression in both the RNA-seq data (Fig. 3e, Supplementary Fig. 7) and in other data sets (Supplementary Fig. 8, 9). Although a small fraction of extreme monoallelic expression is expected to be real, the majority of this excess is likely to result from homozygous individuals that have been miscalled as heterozygotes (Supplementary Fig. 6b). In addition to genotyping errors, RASQUAL can also correct for haplotype switching in heterozygous individuals for rSNPs with large effects (Supplementary Fig. 10). After fitting RASQUAL the frequency of monoallelic expression at heterozygous SNPs was significantly reduced (Fig. 3e). Compared with a model where genotypes and haplotype phase were fixed, the full RASQUAL model exhibited a 9% increase in power (Supplementary Fig. 5), suggesting that our approach significantly improves QTL detection.

### Reference bias and mapping error

AS signals can also be affected by mapping bias towards the reference genome and by mapping errors that assign reads to incorrect genomic locations. Previous approaches, such as the WASP pipeline [6], have used a filtering strategy to remove reads suspected of being influenced by reference bias. In contrast, RASQUAL uses a feature-specific parameter *ϕ* (where *ϕ* = 0.5 denoting no bias towards the reference) to detect individual regions where mapping is biased towards the reference. To test performance of this parameter, we examined the genes for which RASQUAL returned small estimates of 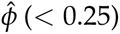. We found that this set was strongly enriched (OR=39.0; *P* = 6.7 *×* 10^*-*22^) for genes located in the MHC region including most known MHC class I & II genes (Fig. 3f). This region is expected to experience severe reference bias because reads overlap multiple alternative alleles in long range LD (Supplementary Fig. 6c). This suggests that RASQUAL efficiently detects and accounts for reference bias without removal of read data.

An additional problem in AS analysis are reads that map to incorrect genomic locations, due to problems in the reference assembly or from sequencing errors. The *d* parameter in RASQUAL captures these mapping errors by comparing genotypes with the observed read sequences during model fitting. We next tested RASQUAL’s ability to capture problems with read mapping in sequenced features. Features exhibiting large *δ* estimates in the RNA-seq data were enriched in pseudogenes (OR=7.6, *P* = 7.5 *×* 10^*-*115^; Supplementary Fig. 12, Supplementary Fig. 6d). They were also significantly enriched in repeat regions and segmental duplications overlapping with CTCF ChIP-seq peaks (OR=3.0, *P <* 10^*-*300^; Supplementary Fig. 13). Additionally, the number of features with large parameter 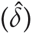 estimates (*>* 1%) was inversely proportional to the length of sequenced read, as would be expected if our parameterisation accurately modelled mapping errors (Supplementary Table 2 and 3).

### Imprinting

An additional feature of RASQUAL is the ability to detect imprinted regions or regions exhibiting RAI. Imprinted regions can be distinguished from *cis*-acting genetic effects because, unlike genetically-driven changes, all samples show allelic imbalance and the identity of the silenced allele is randomised across individuals because maternal and paternal alleles are typically unknown in each individual (Supplementary Fig. 6e). RASQUAL tests for imprinted regions by fitting a model with an imaginary causal variant at which all individuals are heterozygous but the phasing between the causal variant and feature SNPs is unknown (see Supplementary Methods for details). We detected 10 imprinted genes from the RNA-seq data (Fig. 3h, Supplementary Table 3), of which 6 genes were already known (http://www.geneimprint.com/database), a highly significant enrichment (OR=192.3, *P* = 5.0 *×* 10^*-*11^) suggesting RASQUAL can also be utilised to detect novel imprinted genes in cases where multiple heterozygous SNPs are located in the sequenced features. RASQUAL also detected imprinted/RAI regions in ChIP-seq data (Supplementary Table 3) in which three peaks were overlapping within 1kb downstream and upstream of H19 (lincRNA) a known imprinted lincRNA in which the paternal allele is silenced during the early development [30, 31].

### Mapping chromatin accessibility QTLs with RASQUAL and ATAC-seq

We next applied RASQUAL to address a specific biological problem: mapping chromatin accessibility QTLs (caQTLs) in a small sample. We generated genome-wide chromatin accessibility landscapes in 24 LCLs from the 1000 Genomes GBR population using ATAC-seq [18] (see Online Methods and Supplementary Methods for details). Despite the modest sample size RASQUAL detected 2,706 caQTLs at FDR=10% using a permutation test (see Online Methods). Lead SNPs detected by RASQUAL were very highly enriched within the ATAC peak itself (842 peaks; OR=47, *P <* 10^*-*16^) (Fig.4a), with a smaller number in perfect LD with one or more SNPs within the peak (130 in perfect LD with a single fSNP, and 35 with 2 fSNPs). In the set of 1007 lead SNPs within a peak or in perfect LD with an fSNP, the majority (692) overlapped a known transcription factor binding motif that was disrupted by one of the SNP alleles (Fig. 4b). An example caQTL where a putatively causal variant (r2886870) is located within both the ATAC peak and an NF*k*B1 motif is shown in Fig. 4c. This SNP is predicted to produce a large (85%) change in binding affinity, with the predicted change in binding corresponding with a change both in ATAC-seq peak height and in AI at flanking heterozygous fSNPs in individuals that are heterozygous at r2886870.

**Figure 4:**
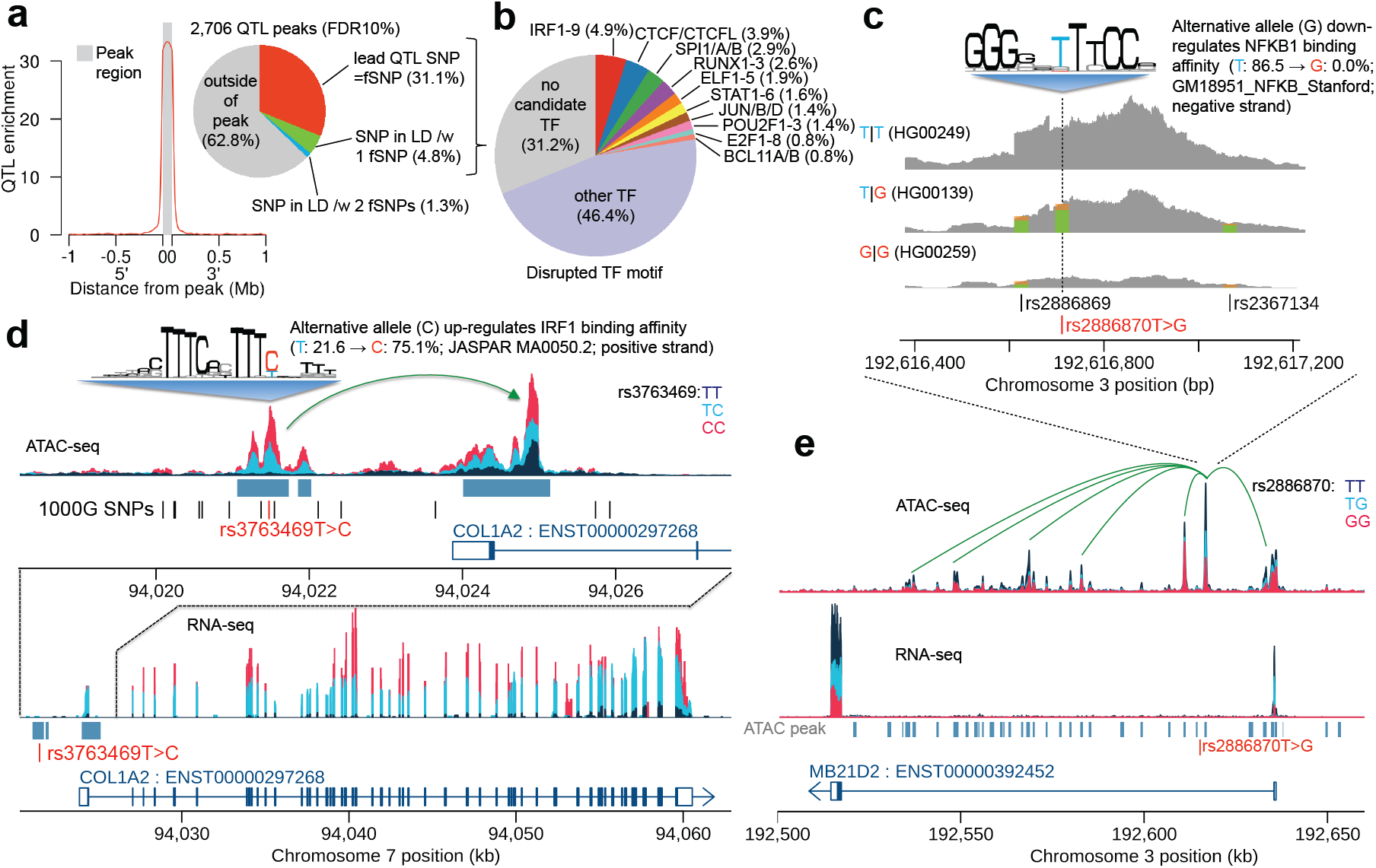
ATAC-QTL mapping with RASQUAL. (a) Positional enrichment of ATAC-QTL lead SNPs, relative to all SNPs, across all 2,706 FDR 10% significant associations detected; inset shows proportion of lead SNPs located inside, outside and in perfect LD (*r*^2^ *>* 0.99) with a SNP inside the ATAC peak. (b) Proportion of lead SNPs located inside, or in perfect LD with a SNP inside the ATAC peak overlapping an identifiable transcription factor binding motif (defined using motifs from the CisBP database [21]). (c) An example of an NF*k*B1 motif-disrupting QTL (d) Example of a “multi-peak” ATAC-QTL (rs3763469) that perturbs a putative enhancer-promoter interaction in the COL1A2, also driving variation in gene expression. Sequence logo illustrates the IRF1 position weight matrix from JASPAR (e) Example of a “multi-peak” QTL: the same genetic variant illustrated in panel c (rs2886870), drives associations at 6 peaks in the intron and promoter the MB21D2 gene.

Further analysis of our detected caQTLs revealed 154 “multipeak” QTLs, where a SNP was associated with variation in chromatin accessibility across multiple independent peak regions. In some cases, these long-range associations appeared to result from enhancer-promoter interactions that are perturbed by a genetic variant. For example, rs3763469 is the lead caQTL SNP for a region of open chromatin located approximately 2.5kb upstream of the promoter of the COL1A2 gene (Fig. 4d) with the alternative allele predicted to increase binding affinity of the transcription factor IRF1. However, we observed that this SNP is also a QTL for the adjacent ATAC peak located over the promoter region of COL1A2 gene, for which no other common SNPs were annotated in the 1000 Genomes database. In other striking examples, we observed genetic associations spanning a large number of additional peaks spread over many tens of kilobases (Fig. 4d). For example the lead caQTL SNP in Fig. 4c also appeared as the lead SNP or SNP in perfect LD with the lead SNP at 5 other peaks in the intron and promoter regions of MB21D2 gene.

### Fine-mapping disease and cell trait associations with RASQUAL and ATAC-seq

Combined with ATAC-seq, RASQUAL becomes a powerful tool for fine-mapping because lead caSNPs are highly enriched in a relatively small genomic space, the ATAC peak itself. caQTLs significantly overlapped GWAS associated SNPs for a range of traits (see Online Methods for details), most significantly rheumatoid arthritis (OR=5.9, 1.9 *×* 10^*-*5^; Fig. 5a). Enabling us to pinpoint putatively causal variants driving disease associations. For example, our analysis highlighted the RA-associated SNP rs909685, which is both a strong caQTL and eQTL for the SYNGR1 gene, as a likely causal variant located within an ATAC peak downstream of the promoter (Fig. 5b). In other cases, our analysis pinpointed instances of multiple, putatively causal variants located within the same ATAC peak. For example we found a suggestive chronic lymphocytic leukemia susceptibility SNP (rs2521269 [32]) in perfect LD with two putatively causal ATAC variants (Fig. 5c). Strikingly, these SNPs may alter the function of an apparently bidirectional promoter also altering the expression of the two adjacent genes, C11ORF21 and TSPAN32 (Supplementary Fig.s14, 15).

**Figure 5:**
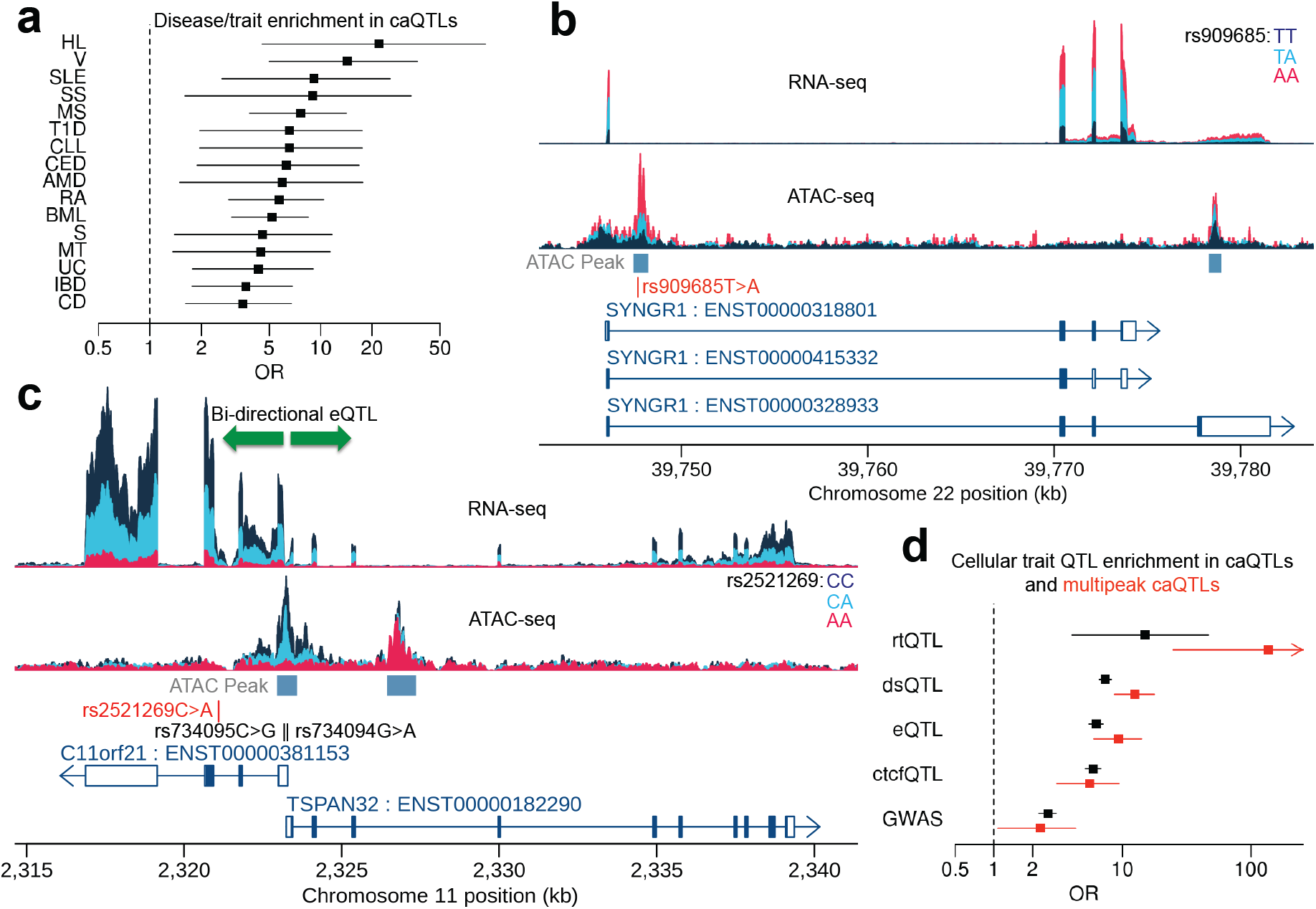
Enrichment of caQTLs and multipeak caQTLs for SNPs associated with other cellular and organismal traits from GWAS. (a) Disease/traits in GWAS catalogue that are enriched in caQTLs (Fisher exact *P <* 0.05). The dot shows the odds ratio between each disease/trait and caQTL, and black line shows its 95% confidence interval. (b) Example of caQTL (rs909685 [40]) that is also RA susceptibility SNP and eQTL for SYNGR1 gene. (c) Suggestive CLL susceptibility SNP (rs2521269 [32]) is a joint ATAC-expression QTL. The alternative allele down-regulates chromatin accessibility and expression levels of two flanking genes (C11orf21 and TSPAN32) simultaneously. The SNP is in perfect LD with two adjacent fSNPs, rs734095 and rs734094, in the ATAC peak (see Supplementary Figure 14 and 15 for details). (d) Cellular trait QTL enrichment in caQTL (black) and multipeak caQTL (red). The dot shows the odds ratio between each disease/trait and caQTL, and the black line shows its 95% confidence interval. The red arrow shows the confidence interval continues toward 582. Hodgkin’s lymphoma (HL); Vitiligo (V); Systemic lupus erythematosus (SLE); Systemic sclerosis (SS); Multiple sclerosis (MS); Type 1 diabetes (T1D); Chronic lymphocytic leukemia (CLL); Celiac disease (CED); Age-related macular degeneration (AMD); Rheumatoid arthritis (RA); Blood metabolite levels (BML); Schizophrenia (S); Metabolic traits (MT); Ulcerative colitis (UC); Inflammatory bowel disease (IBD); Crohn’s disease (CD); DNA replication timing QTL (rtQTL); DNaseI hypersensitive QTL (dsQTL); CTCF binding QTL (ctcfQTL); expression QTL (eQTL).

The caQTLs we detected were also significantly enriched for other cellular QTLs detected in LCLs including DNaseI-seq, CTCF ChIP-seq and RNA-seq data sets (Fig. 5d), with multipeak QTLs more than twice as likely to be associated with gene expression than normal caQTLs. Strikingly, our caQTLs were most strongly enriched in replication timing QTLs (rtQTLs) (OR=15.4; *P* = 3.0 *×* 10^*-*4^) recently mapped in LCLs [33]. This enrichment was even more extreme when we considered multipeak caQTLs, which were 10 times more likely to be associated (OR=159.3, *P* = 1.8 *×* 10^*-*6^; Fig. 5d) than normal caQTLs. The example multipeak QTL SNP rs2886870 (Fig. 4e) is in perfect LD with the rtQTL SNP (rs6786283) detected in ref [33] in Europeans.

## Discussion

We have developed a novel statistical model, RASQUAL, for mapping associations between genotype and NGS-based cellular phenotypes. In our tests, RASQUAL consistently outperformed all other existing methods across a range of NGS data types. We generated a novel ATAC-seq data set in LCLs from European individuals and illustrated how RASQUAL can be used for fine-mapping disease-associated variants and for uncovering fundamental mechanisms of gene regulation.

A major difference between RASQUAL and the other methods we have tested is that RASQUAL handles bias and detection of genetic signals in a single statistical framework, using information from all individuals in the data set and without removing data. In contrast, other methods treat bias in NGS data as a data quality control (QC) issue and either ignore it or rely on a series of read realignment and data filtering steps to remove problematic regions *a priori*. Better handling of a range of biases in NGS data is likely to explain much of the differences in performance we observed. Furthermore, data filtering and QC steps as an alternative to handling bias may introduce a number of other issues. First, data QC involving read filtering may also inadvertently remove substantial signal from the data. As one example, we found that the WASP pipeline removed between 22 and 31% of reads in our RNA-seq subset analysis, but that this appeared to make a relatively minor difference to power for association detection (Supplementary Fig. 11a). A second issue is that it is often difficult to set sensible thresholds for data QC. In addition to boosting model sensitivity and specificity, we believe that minimising the amount of data pre-processing significantly improves the usability of RASQUAL. Users of RASQUAL are not required set arbitrary thresholds for data QC, nor are they required to perform disk and CPU-intensive read remapping or simulations to account for biases NGS data, but can instead run the model on existing data as is. In addition, RASQUAL output contains an informative set of parameters that can highlight genomic regions with problematic AS signals enabling more informed downstream interpretation of analysis results.

A potential limitation of RASQUAL is its reliance on genotype imputation and phasing to compute genotype and diplotype likelihoods. Poor quality imputation or phasing information is likely to affect the RASQUAL’s ability to detect QTLs, particularly in cases where the distance between the true rSNP and fSNP is large, due to the increased probability of haplotype switching errors. However our analysis illustrates that these problems can be readily overcome using imputation into the large, high quality panels now available from the 1000 Genomes and UK10K projects. Additionally, RASQUAL can be run in a “genotype-free” mode, where only SNPs located inside sequenced features are considered, genotypes are learned from the read data and SNP locations are specified using, for example, dbSNP. Although lack of genotype information will reduce power substantially, it can enable analysis of NGS data sets where genotype data are lacking [34].

Our results also illustrate how RASQUAL can be used to extract meaningful genetic signals from data sets of a modest size. For example, our analysis of ATAC-seq data demonstrates how genetic variation can be leveraged to connect distal regulatory elements with gene promoters or with other regulatory elements. A strength of this approach, compared with experimental techniques such as Hi-C or CHiAPET, is that these interactions are linked to specific genetic changes enabling characterisation of causal relationships between regulatory elements and their target genes. We expect that genetic analysis of long-range regulatory interactions will be a powerful complement to standard experimental techniques in future studies. Our analysis also uncovered a potential link between replication timing and chromatin openness. A possible explanation for this result is that between-individual variation in replication timing manifests as a change in DNA copy number that, in turn, appears as a change in open chromatin. However, the number of cells in S-phase is likely to be between 5-20% at any given time [33] implying an absolute maximum expected fold change of 1.2 (assuming all individuals have 20% of cells in S-phase at the same time). This effect seems insufficient to explain the magnitude of ATAC QTL effect we observe in many cases with, for example, a 2.3-fold change in ATAC-seq peak height between the major and minor homozygotes in Fig. 4e.

Finally, we believe that our results also highlight how RASQUAL’s performance with modest sample sizes will enable researchers to collect and analyse multiple complementary NGS data sets, rather than focusing resources on maximising sample sizes for an individual phenotype. Combined with RASQUAL’s improved ability to localise causal variants we suggest that a major future application of our model will be the fine-mapping of causal regulatory variants to understand the molecular mechanisms underlying phenotypic variation.

## Online methods

### Statistical overview of RASQUAL

RASQUAL models each sequenced feature, such as ChIP-seq peak or the union of exons over an entire transcript, and considers all genotyped variants within a given distance of the feature (the *cis*-window). For simplicity, RASQUAL assumes a single causal variant at each feature, although multiple causal variants can be tested for by conditioning on the lead SNP genotype. Let *Y*_*i*_ be the total fragment count at the feature for individual *i* (*i* = 1, …, *N*), ***G***_*i*_ be the putative causal genetic variant in the region and ***G***_*il*_ be the *l*-th variant within the sequenced feature, such that we observe allele-specific fragment counts 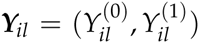 at each feature variant *l* (Fig. 1).

The model contains two components: (i) population signals are captured by regressing the total fragment count *Y*_*i*_ onto the number of alternative alleles *G*_*i*_ (*G*_*i*_ = 0, 1, 2) at the putative cis-regulatory SNP, assuming read counts follow a negative binomial distribution (*p*_NB_) and (ii) AS signals are modelled assuming the alternative fragment count at a fSNP *l* given the total reads overlapping that fSNP follows a beta binomial distribution (*p*_BB_). The model components are connected by a single cis-regulatory effect parameter *π* such that the expected total count is proportional to {2(1 *π*), 1, 2*π*} for *G*_*i*_ = 0, 1, 2 and the expected allelic ratio in an individual heterozygous for the putative causal SNP becomes *{π*,1 *-π*} as in [6, 19].

The likelihood is then given by

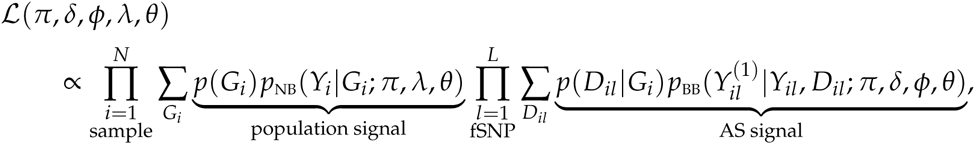

where *D*_*il*_ denotes the diplotype configuration in individual *i* between the putatively causal variant and the fSNP *l*, *p*(*G*_*i*_) and *p*(*D*_*il*_*|G*_*i*_) denote prior probabilities of genotype and diplotype configuration (obtained from SNP phasing and imputation). In addition to the *cis* genetic effect (*π*), total read count depends on *δ*, a scaling parameter for absolute mean of the negative binomial distribution. The allelic ratio depends upon *ϕ*, the probability that an individual read maps to an incorrect location in genome and *δ*, the reference mapping bias (where *ϕ*=0.5 corresponds to no reference bias). Overdispersion in both *Y*_*i*_ and 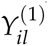 is captured by a single shared parameter *θ*.

For statistical hypothesis testing of QTL, all five parameters for each SNP-feature combination in the *cis*-regulatory window are estimated independently to get the maximum likelihood under alternative hypotheses. Under the null hypothesis, all parameters except *π* are estimated for each feature independently, while *π* is set to 0.5 and we use a likelihood ratio test to compare the null and alternative hypotheses for each SNP-feature combination using the *χ* square distribution with one degree of freedom (for *π*). We use an EM algorithm to obtain the maximum likelihood estimators of the parameters [20]. We do not introduce any common parameters across individuals or features estimated *a priori*, but instead introduced prior distributions for all the parameters (see Supplementary Methods for details) to increase the stability and usability of RASQUAL. A detailed description of the derivation of statistical model and the EM algorithm is available in the Supplementary Methods.

### Data preprocessing of sequencing traits

The GEUVADIS RNA-seq data was downloaded from ArrayExpress (Accession E-GEUV-3), CTCF CHIP-seq data was downloaded from the European Nucleotide Archive (Accession ERP002168) and the DNase1-seq was downloaded from the Gene Expression Omnibus (Accession GSE31388). All data sets were realigned to human genome assembly GRC37. RNA-seq data were aligned using Bowtie 2 [35] and reads mapped to splice junctions using tophat 2 [36], with ENSEMBL human gene assembly 69 as the reference transcriptome. CTCF ChIP-seq data was realigned using bwa [37] and the DNase1-seq was realigned using the alignment method described in Degner *et al* [5]. Following alignment, we removed reads with a quality score of *<*10 from all three data sets.

For the CTCF ChIP-seq and DNase1-seq data, we generated genome wide read coverage depths from either the fragment midpoints or cut site data respectively. Peaks were called by comparing two Gaussian kernel densities with bandwidths of 100 and 1000bp, corresponding to a “peak” and “background” model respectively. We then defined a peak as a region where the peak kernel coverage exceeded the background kernel coverage, and where the peak coverage was greater than 0.001 fragments per million.

For RNA-seq data, we counted the number of sequenced fragments (mate-pairs) of which one or other sequenced end overlaps with an union of annotated Ensembl gene exons. For CTCF ChIP-seq and ATAC-seq data, we counted the number of sequenced fragments of which one or other sequenced end overlaps with the annotated peak. For DNase-seq data, we simply counted the number of reads that are overlapping with the annotated peak. For the computation of principal components we also calculated FPKM and RPKM values for these data sets (see Supplementary Methods). All sequence data sets were corrected for between library variation in amplification efficiencies of different GC content reads. For each sample, all features were binned based on their GC content, the relative over-representation of features of a given GC content for a given sample relative to all other samples was estimated using a smoothing spline. This value was then either included as a covariate, in the comparison of CHT, TReCASE and RASQUAL, or to correct RPKM or FPKM values for the linear model.

### SNP genotype data preparation

We downloaded VCF files for the 1000 Genomes Phase I integrated variant set from the project website. Because RNA-seq and ATAC-seq samples completely overlapped with the 1000 Genomes samples, we used the subsamples from the VCF files. For CTCF ChIP-seq and DNaseI-seq data, samples were completely overlapped with the HapMap samples (except for NA12414 in CEU population and NA18907 in YRI population) but not 1000 Genomes samples. Therefore we downloaded the HapMap phase II & III genotypes from the project website and imputed with the 1000 Genomes Phase I haplotypes using IMPUTE2 [38]. For the two samples which are not in HapMap samples, we obtained genotypes from the 1000 Genomes data at HapMap SNP loci and merged before the imputation. We adopted the common 2-step imputation approach to phase HapMap genotypes first and then impute haplotypes. Note that, to apply whole genome imputation, we split each chromosome in 20Mb bins with 100Kb overlaps.

For any cellular trait mapping, we used SNP loci with minor allele frequency greater than 5% and imputation quality score (MaCH *R*^2^ or IMPUTE2 *I*^2^) greater than 0.7 for candidate rSNP. For fSNPs, we used all SNPs overlapping with the target feature with at least one individual being heterozygote. For TReCASE analysis, we merged AS counts at those fSNPs with heterozygous genotypes for each feature according to the phased haplotype information. Note that, indels and other structural variants were discarded.

### Definition of true QTLs

We downloaded the eQTL list detected using the entire European data set (*N* = 373) at FDR 5% from (http://www.ebi.ac.uk/arrayexpress/files/E-GEUV-1/analysis_results/EUR373.gene.cis.FDR5.best.rs137.txt.gz) as a gold standard eQTL gene set. We also obtained the list of dsQTL regions detected using the 70 YRI individuals under FDR 10% (http://eqtl.uchicago.edu/dsQTL_data/QTLs/GSE31388_dsQtlTable.txt.gz). and used the liftover tool to (http://genome.sph.umich.edu/wiki/LiftOver) to transfer genome coordinates on hg19. We then obtained peaks (in our annotation) that overlapped with the reported dsQTL regions as a gold standard dsQTL peak set.

### CTCF motif-disrupting SNPs

Motif-disrupting SNPs were defined as SNPs located within a CTCF peak and putative CTCF motif, whose predicted allelic effect on binding (computed using CisBP [21] position weight matrices (PWMs) corresponded to an observed change in CTCF ChIP-seq peak height in the expected direction.

The predicted allelic effect is calculated from a PWM as follows. Let *S*_*a*:*b*_ be the reference sequence at chromosomal position between *a* and *b* on a chromosome. We assume a SNP locus at chromosomal position *c*. For a PWM with motif length *m*, we calculate the binding affinity score

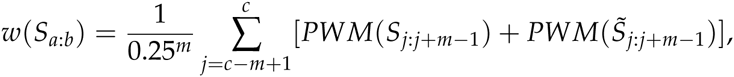

where *PWM*(*·*) denotes the PWM score for *S*_*a*:*b*_ and 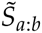 denotes the reverse complement sequence of *S*_*a*:*b*_. We also calculated the affinity score for the sequence 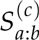 where the reference sequence at position *c*, that is *S*_*c*:*c*_, is replaced by the alternative allele of the SNP. We compared *w*(*S*_*a*:*b*_) with 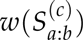 to determine which SNP allele is over-represented at the putative binding site involving the SNP locus at *c*.

For CTCF-binding motifs, there exist multiple PWMs (*N* = 67) reported in [21]. We simply took the average affinity score 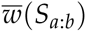 across all PWMs. Then we considered only SNPs that gives either 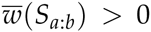 or 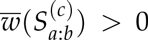 as a SNP in a CTCF motif starting at chromosomal position 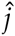, such that

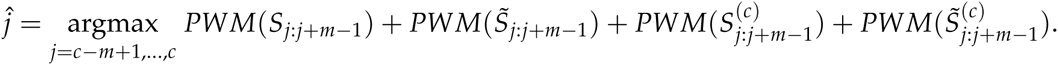

### Multiple testing correction

We implemented a two-stage multiple testing correction to determine which features is a significant QTL. First, because SNP density varies between genomic regions QTL mapping for different features involves testing different number of SNPs. This results in lead *P*-values that are incomparable across features because more SNP dense regions will involve greater numbers of tests and therefore smaller *P*-values observed by chance under the null hypothesis. Second, after *P*-values for each feature have been corrected for the number of tests in the *cis*- window, they are corrected again for the number of features tested genome-wide. In our analysis we used a multiple-testing correction that used the Beta distribution for correction within each *cis*-window, followed by a permutation test that corrects for the number features tested genome-wide.

Previous studies have used a Bonferroni correction to correct for the multiple testing within *cis*-windows [4]. However, the Bonferroni correction becomes very aggressive when sample sizes are small and asymptotic normality does not hold. In such cases, *P*-values tend to become inflated and skewed towards 1. We observed that the Bonferroni method aggressively corrected for *P*-values with greater numbers of tests, and frequently significant QTLs were only detected for features where the number of tests was relatively small (see Supplementary Methods and Supplementary Fig. 17) while QQ-plots revealed that Bonferroni corrected *P*- values were extremely skewed towards 1 (Supplementary Fig. 17). To deal with this problem, we instead assumed *P*-values were drawn from a Beta distribution with scale parameters (*α*, instead of the uniform distribution (*α*=*β*=1) assumed in the Bonferroni method. We fit the distribution derived from the first order statistic of a series of Beta random variables to correct for the multiple testing (see Supplementary Methods for details). The Beta correction for multiple testing resulted in *P*-values that were much better calibrated in small samples (Supplementary Fig. 17d) and so we consistently used this multiple testing correction for *P*-values from RASQUAL and also other methods (CHT, TReCASE and linear model). We note that the major conclusions of our manuscript were not affected by the choice of multiple testing correction, with RASQUAL outperforming TReCASE and CHT regardless of which multiple testing correction we used (Supplementary Fig. 18).

After *P*-values within each feature were corrected, we again corrected for the number of features tested genome wide. Although *P*-values were well calibrated after the Beta correction, they can still deviate from uniformity in smaller sample sizes. To account for this, we used a permutation test. Here, we draw random permutations {(*i*)} for total fragment count and {(*i*_*l*_)} for AS counts at each fSNP *l* independently. Then we maximise the following likelihood

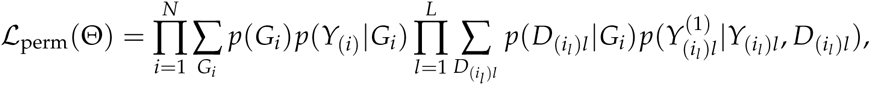

with respect to θ = (*π*, *δ*, *ϕ*, *λ*, *θ*) to obtain the likelihood ratio statistic (for *π* = 0 and *π ≠ 0*) under the putative null hypothesis. Here, *D*_(i_l_)l_ denotes the diplotype configuration between *G*_*i*_ and permuted fSNP *G*_(i_l_)l_. *P*-values obtained from permuted data were corrected for multiple tests within each feature as described for real data. Then the permutation *P*-values were compared with the real *P*-values to calibrate genome-wide false-discovery rate. Note that the permutation *P*-values can also be used to estimate the scale parameters (*α*, *β*) in the Beta correction to simulate a more realistic null distribution. For caQTL mapping, we used those scale parameters estimated from the permutation test for the read *P*-values to correct for the multiple testing within each feature.

#### ATAC-seq in LCLs

The ATAC-seq method used was as described in [18], but with some modifications: (1) 100,000 LCL nuclei obtained from sucrose and Triton X-100 treatment were tagmented using the Illumina Nextera kit and then subject to limited PCR amplification, incorporating indexing sequence tags (2) ATAC libraries were purified and size selected before pooling (3) index tag ratios were balanced using a MiSeq (Illumina) run before deep sequencing with 75bp pairedend reads on a HiSeq 2500 (Illumina). For more details, see Supplementary Methods.

#### Disease enrichment analysis for ATAC QTLs

We obtained the publicly available GWAS catalogue data [39] from the UCSC website created on Mar 2015. We only included studies that had at least 10 hits that were genome-wide significant of *P <* 5 *×* 10^*-*8^, that overlapped with the SNPs tested in ATAC QTL mapping (5,703,168 loci as a total) and were based on European populations with the sample sizes greater than 1000. The resulting data set contained GWAS on 101 diseases and other traits. Because of tight LD, different index SNPs in the same locus were reported by multiple GWA studies for a single disease/trait. Likewise, multiple LD SNPs were significantly associated with a single ATAC peak. To merge these LD SNPs, we assigned the lead ATAC peak with the minimum *P*-value for each SNP locus and counted the number of lead peaks (instead of SNPs) that are significantly associated with a disease/trait and/or ATAC QTLs (Supplementary Fig. 16). The disease/trait enrichment was assessed using a Fisher exact test. Note that, the number of tested peaks is different across SNPs because multiple testing correction has been applied for each lead *P*-value and SNPs with the corrected lead *P*-values less than FDR10% were called as significant ATAC QTL SNPs.

